# Emergent *emm4* group A *Streptococcus* evidences a survival strategy during interaction with immune effector cells

**DOI:** 10.1101/2024.04.09.588776

**Authors:** Chioma M. Odo, Luis A Vega, Piyali Mukherjee, Sruti DebRoy, Anthony R. Flores, Samuel A. Shelburne

## Abstract

The major gram-positive pathogen group A *Streptococcus* (GAS) is a model organism for studying microbial epidemics as it causes waves of infections. Since 1980, several GAS epidemics have been ascribed to the emergence of clones producing increased amounts of key virulence factors such as streptolysin O (SLO). Herein, we sought to identify mechanisms underlying our recently identified temporal clonal emergence amongst *emm4* GAS, given that emergent strains did not produce augmented levels of virulence factors relative to historic isolates. Through the creation and analysis of isoallelic strains, we determined that a conserved mutation in a previously undescribed gene encoding a putative carbonic anhydrase was responsible for the defective *in vitro* growth observed in the emergent strains. We also identified that the emergent strains survived better inside macrophages and killed macrophages at lower rates relative to the historic strains. Via creation of isogenic mutant strains, we linked the emergent strain “survival” phenotype to the downregulation of the SLO encoding gene and upregulation of the *msrAB* operon which encodes proteins involved in defense against extracellular oxidative stress. Our findings are in accord with recent surveillance studies which found high ratio of mucosal (i.e., pharyngeal) relative to invasive infections amongst *emm4* GAS. Inasmuch as ever-increasing virulence is unlikely to be evolutionary advantageous for a microbial pathogen, our data furthers understanding of the well described oscillating patterns of virulent GAS infections by demonstrating mechanisms by which emergent strains adapt a “survival” strategy to outcompete previously circulating isolates.

## INTRODUCTION

Microbial pandemics/epidemics are major drivers of human disease and can result from the acquisition of novel genetic material or mutation of existing material which confers a fitness advantage [1–4]. For example, the signature mutations acquired by the highly transmissible Omicron variant of SARS-CoV-2 have resulted in decreased immune control in persons with previous COVID-19 as well as thwarted immunization efforts [5–8]. With the increasing availability of whole genome sequencing of large cohorts of microbial pathogens, it has become apparent that emerging strains are often separated from precursors by a small number of genetic differences which provide opportunities to dissect molecular mechanisms driving clonal epidemics [9–12].

Group A *Streptococcus* (GAS) is a model for studying molecular pathogenomics of microbial epidemics as it has a relatively small genome (∼2.0 Mb), there is active surveillance for GAS strains in many countries, and there is significant serological diversity (more than 200 *emm* types) based on the 5’ sequence of the *emm* gene encoding the N-terminal region of the surface M protein [13–16]. GAS epidemiology has been characterized by large fluctuations in disease incidence and severity, such as the near disappearance of GAS-induced rheumatic fever from North America and Western Europe in the latter half of the 20th century [17–19]. The molecular basis for such variances remains incompletely understood, although recent advances have been made using systematic, whole-genome sequencing (WGS) efforts [20–22]. For example, the high lethality of currently circulating *emm3* GAS strains has been linked to the acquisition of an actively secreted phospholipase [23]. Similarly, the recent emergence of hypervirulent *emm1* and *emm89* GAS clones has been attributed to recombination in the *nga/slo* promoter leading to augmented production of the Nga and Streptolysin O (Slo) toxins, which act synergistically to kill human cells [17, 20, 24, 25]. Finally, recent upsurges in scarlet fever have been attributed to a single polymorphism in the 5’ UTR of *speA* in *emm1* strains which increases production of the superantigen SpeA [26–28].

A clonal emergence, expansion, and replacement involving *emm4 GAS* strains in the United States and the United Kingdom was recently identified [29, 30]. Using invasive strains collected as part of active surveillance, we estimated that a new *emm4* clone emerged around 1996 and, by 2017, had completely replaced the existing historic *emm4* strains [31]. There were a limited number of genetic changes consistently separating the new “emergent” from the old “historic” *emm4* strains [31]. These included a gene fusion between the *emm* and adjacent *enn* genes resulting in a chimeric M protein, a frameshift mutation in *silA* [32, 33], a premature stop codon in the gene encoding the regulatory protein Ralp3 [34], and early termination in the gene encoding a putative carbonic anhydrase enzyme. Surprisingly, the emergent strains had lower transcript levels and protein production of *nga/slo* and had not acquired a new virulence factor, suggesting a possible novel mechanism underlying GAS strain emergence [31]. Thus, we sought to gain insight into the mechanisms driving strain replacement in *emm4* GAS through a detailed examination of the genotypic and phenotypic differences between the historic and emergent *emm4* strains.

## RESULTS

### Analysis of the role of *ralp3* and *silA* mutations in *emm4* GAS

In light of the significant differences in the transcriptomes of the historic and emergent *emm4* strains [31], we first focused our attention on the mutations in genes encoding regulatory proteins. All of the emergent strains have an early stop codon predicted to lead to a truncated Ralp3 [31], a transcriptional regulator only present in some GAS *emm* types which has been previously shown to contribute to the virulence of *emm1* GAS [34, 35]. To evaluate the impact of the premature *ralp3* stop codon, an emergent strain with a full-length Ralp3 was constructed (ABC208 *ralp3^WT^*) using an isoallelic strategy [36]. Given that Ralp3 is reported to regulate virulence factors in *emm1* and *emm49* GAS [34, 35], we then evaluated the impact of Ralp3 restoration on the transcript levels of some of the genes previously found to be influenced by Ralp3 such as *mga*, which encodes a critical virulence factor regulator [37] and the cysteine protease encoding gene *speB* [38]. Although there were differences in transcript levels between the historic wildtype (WT) (ABC25) and emergent WT strains (ABC208), fixing the *ralp3* mutation in the emergent strain did not significantly impact mga or *speB* transcript levels (*P* > 0.05) (Fig. 1A-B). In light of previous data noting the importance of Ralp3 on GAS survival in human blood [34], we assayed the impact of the Ralp3 restoration on GAS-macrophage interaction. Similar to the transcript level data, there were survival differences between the historic and emergent WT strains, but there was no significant impact of the Ralp3 restoration on this phenotype (*P* > 0.05) (Fig. 1C). Of note, both the historic and emergent *emm4* strains contain the unusual ATA start codon for *ralp3* as opposed to the standard ATG for *emm1* GAS suggesting that the mRNA may not be translated into a significant amount of protein even when the gene is intact [39].

**FIG 1.**
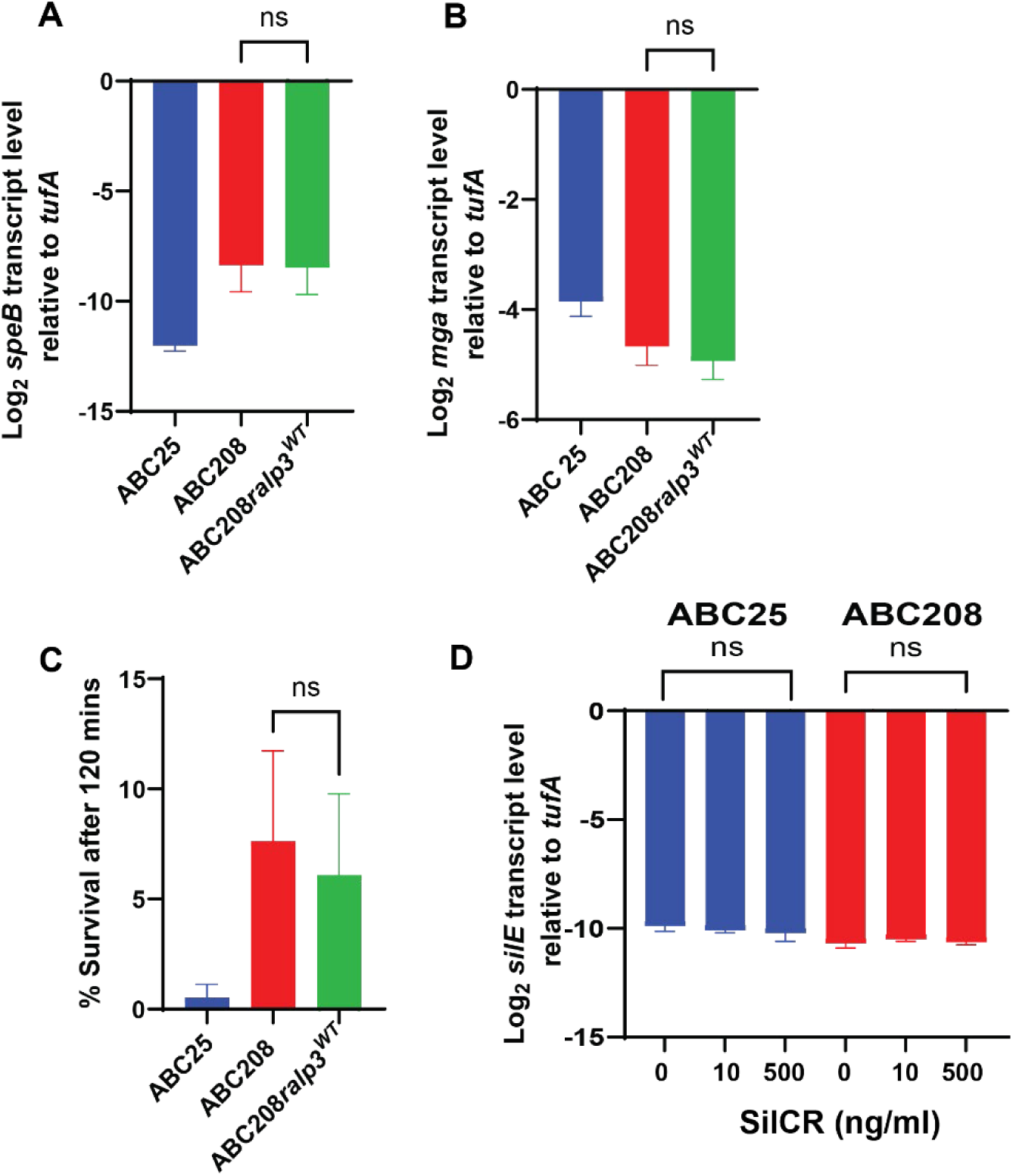
Analysis of the role of *ralp3* and *silA* mutations in *emm4* GAS. (A-B) Log_2_ transcript levels (qRT-PCR) of *speB* and *mga* from ABC25 (historic strain in blue), ABC208 (emergent strain in red), and ABC208*ralp3^WT^* (in green). (C) Survival of indicated strains after interaction with J774A.1 macrophages for 120 mins. (D) Log_2_ transcript level (qRT-PCR) of *silE* from ABC25 and ABC208 grown in different concentrations of SilCR peptide as indicated on the x-axis. For each panel, the error bars represent the standard deviation of three biological and two technical replicates. ns = no significant difference by either Student’s t-test (panels A-C) or ANOVA (panel D).

The *silA* gene encodes the response regulator portion of the *Streptococcus* invasion locus (Sil) quorum sensing system, which is present in a limited number of GAS *emm* types and is important in necrotizing fasciitis [40, 41]. Whereas frameshift of the *silA* gene is only present in emergent *emm4* GAS, both emergent and historic strains contain an early stop codon in the *silD* gene, which encodes an ABC transporter, raising the possibility that the system is not active in either the historic or emergent strains [42]. We tested the functionality of the *sil* system in both the historic strain ABC25 and emergent strain ABC208 by growing the strains in different concentrations of the activating peptide SilCR and analyzing transcript levels of *silE*, which markedly increases in the presence of SilCR when the system is intact [32, 43]. Consistent with a non-functional Sil system, we found no significant impact on *silE* transcript levels at any concentration of SilCR (*P* > 0.05 by ANOVA) (Fig. 1D). Taken together, we conclude that is unlikely that the mutations in *ralp3* or *silA* markedly impact the pathophysiology of emergent *emm4* GAS.

### Creation and evaluation of a historic *emm4* GAS strain producing a chimeric M/Enn protein

Considering the well-established role of M protein as a key, cell-surface anti-phagocytic virulence factor [44–46], we sought to assess the impact of the *emm-enn* gene fusion event by constructing a historic strain with an emergent chimeric *emm-enn* gene, herein called ABC25-chimera (Fig. 2A). Additionally, we created an *emm*-knockout in the representative emergent strain ABC208 (ABC208Δ*emm*). As genetic manipulation of the *emm* region can result in the silencing of *mga* and *emm* transcription, we took the following steps to ensure that our strain creation strategy did not lead to “off-target” effects [47]. First, we performed Sanger and whole genome sequencing to verify accuracy of the genetic constructs and lack of secondary mutations respectively. Second, we analyzed the *emm* transcript levels in all strains. Finally, we ensured that M protein was being produced and present on the cell surface at similar levels between parent and derivative strains using both Western immunoblots of whole cell lysates as well as cell-surface analysis via flow cytometry, respectively.

**FIG 2.**
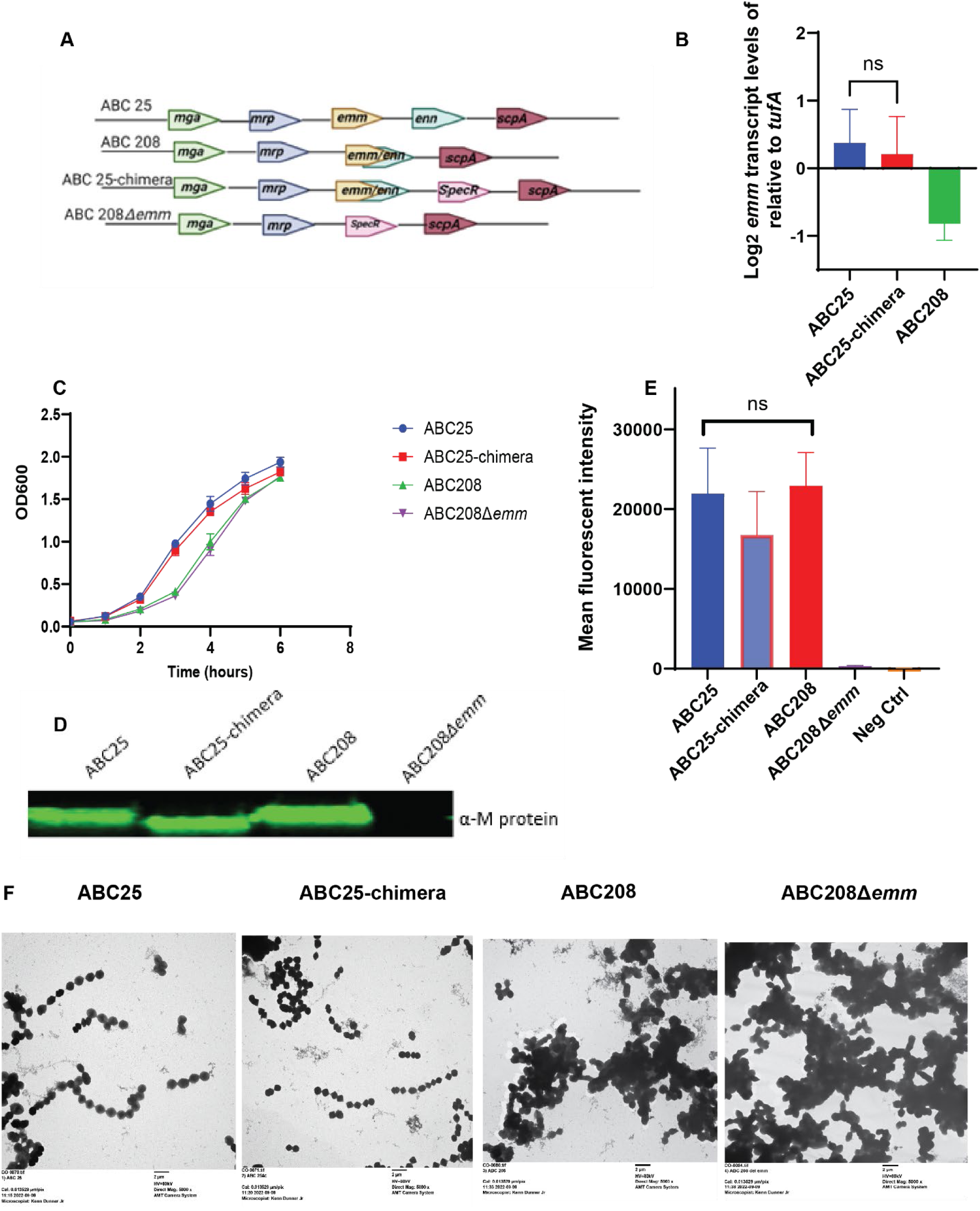
Characterization of *emm-enn* gene fusion in *emm4* GAS. (A) Arrangement of *emm* and the surrounding genes in the *mga* locus in clinical and genetically engineered *emm4* strains. (B) Growth curves in THY for ABC25 (historic), ABC208 (emergent) ABC25-chimera (historic with emergent M protein) and ABC208Δ*emm* (emergent without *emm* gene). (C) *emm* transcript levels from indicated strains grown in THY to ME phase. Error bars represent the standard deviation of three biological and two technical replicates. (D) Western blot analysis using anti-M4 antibody (α-M) of whole cell lysate from indicated strains. Note that M protein in ABC-25 chimera runs slightly smaller relative to parent as it lacks a 35 amino acid C-repeat [29]. (E) Flow cytometry analysis of surface localized M protein on indicated strains using anti-M4 protein antibody. NS indicates *P* > 0.05 by either Student’s t-test (panel B) or ANOVA (panel E). (F) Transmission electron microscopy images of indicated strains (Mag-5000 x).

As expected, in THY the historic strain ABC25 grew better relative to the emergent strain ABC208, but we observed no growth impact for either varying or removing the *emm* gene relative to the respective counterparts (Fig. 2B). The qRT-PCR analysis showed slightly lower *emm* transcript levels in ABC208 relative to ABC25, but no there was significant difference in *emm* transcript levels between ABC25 and the ABC25-chimera (*P* > 0.05) (Fig. 2C). The western blot analysis of whole cell lysates showed similar M protein levels in ABC25, ABC25-chimera, and ABC208 with no detectable signal in ABC208Δ*emm* (Fig. 2D). By flow cytometry using a custom-made anti-M4 antibody, there was no significant difference in mean fluorescent intensity between ABC25, ABC208, and ABC25-chimera (*P* > 0.05 by ANOVA), whereas the mean fluorescent intensity in ABC208Δ*emm* was similar to that of the negative control (Fig. 2E). From these data, we conclude that the genetic creation of a historic strain with a chimeric *emm-enn* gene did not significantly impact *emm* transcript levels or M protein production/cell surface localization.

We began to evaluate the impact of the *emm-enn* gene fusion by studying strain morphology given that M protein has previously been identified as an important contributor to GAS chain length [48]. To this end, strains were analyzed using transmission electron microscopy (TEM), and we found that both ABC25 and ABC25-chimera had typical streptococcal chains whereas strains ABC208 and ABC208Δ*emm* were characterized by clumping cells (Fig. 2F). Taken together, these data show that whereas there are growth and morphological differences between historic and emergent *emm4* strains [31], these differences are not driven by the *emm-enn* gene fusion.

### Identification of carbonic anhydrase as critical to differences in growth between emergent and historic *emm4* GAS

Carbonic anhydrase catalyzes the reversible hydration of CO_2_ to aqueous bicarbonate (HCO_3_) and is essential for *in vitro* growth in *Streptococcus pneumonia* [49] but has not been characterized in GAS. We identified an early stop codon in the carbonic anhydrase encoding gene (herein named *saca* for *<u>S</u>treptococcus* <u>A</u> <u>c</u>arbonic <u>a</u>nhydrase) only in the emergent *emm4* strains [31]. To determine whether this variation impacted previously observed differences in growth between the emergent and historic strains (Fig. 2B) [31], we constructed an emergent strain with a WT *saca* gene (ABC208*saca^WT^)* and a historic strain with an early stop codon in the *saca* gene (ABC25*sacaQ12*)*. We found that repairing the mutation in the *saca* gene in ABC208 resulted in a similar THY growth phenotype to ABC25 whereas introducing the mutation into ABC25 led to a marked growth defect (Fig. 3A). Also, growing ABC208 in the presence of HCO_3_ markedly improved its growth to a level similar to that of ABC25 (Fig. 3B). These data indicate that the early stop codon in *saca* is a major contributor to the in vitro growth characteristics separating historic from emergent *emm4* GAS.

**FIG 3.**
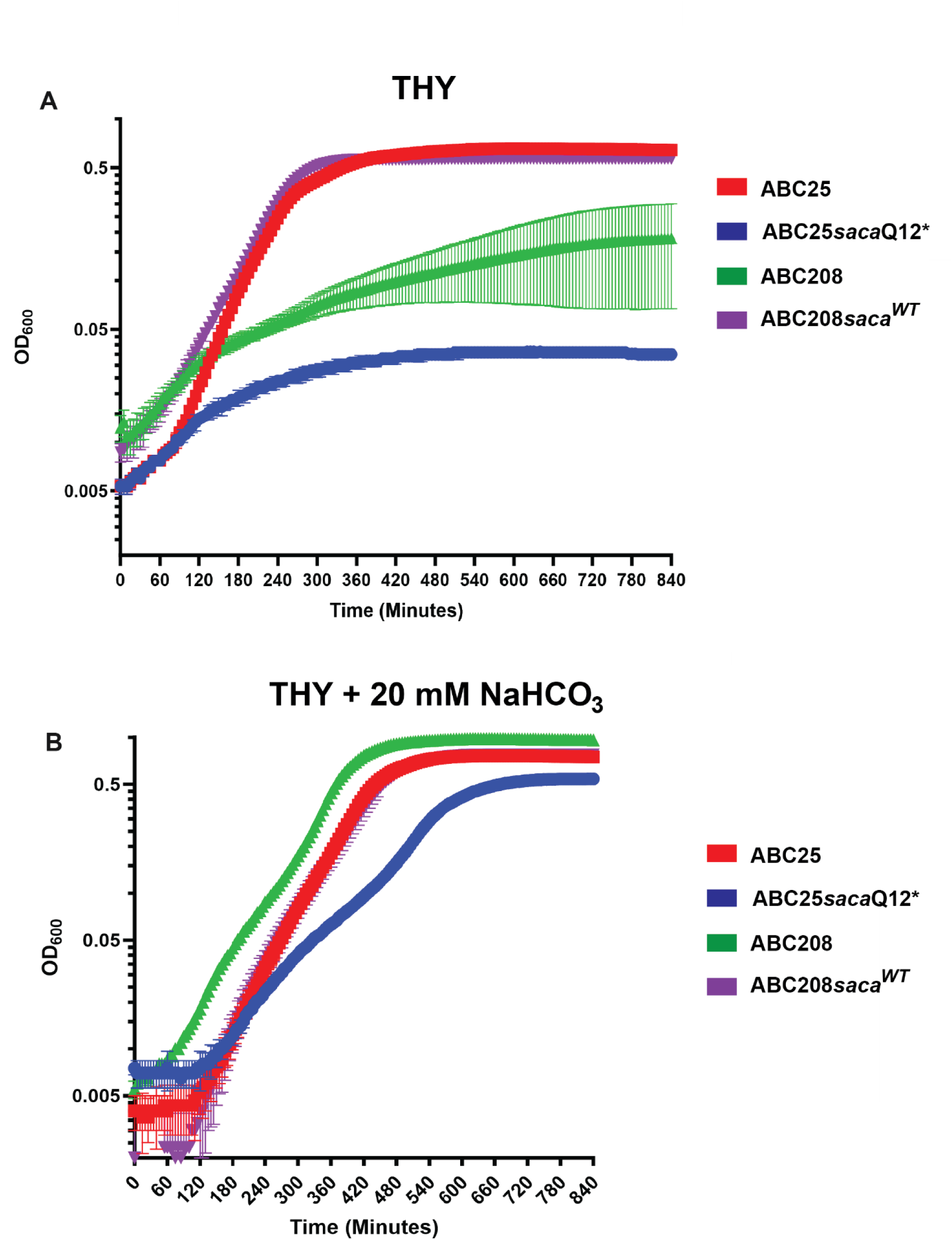
Characterization of the impact of mutation in a putative carbonic anhydrase encoding gene on *emm4* GAS growth. (A) THY growth analysis of the historic strain ABC25 (blue), ABC25 *saca* mutant (green), the emergent strain ABC208 (red), and ABC208 with restored *saca* gene (purple). (B) Same strains grown in THY supplemented with 20mM NaHCO_3_. For both (A) and (B), error bars represent the standard deviations for four replicates per strain on three separate days.

### Assessment of binding of fibrinogen and C4 binding protein between historic and emergent strains

The GAS M and M-related proteins have diverse interactions with the human immune system including binding to fibrinogen and C4 binding protein (C4BP) [50–53]. Given that the *emm-enn* gene fusion could, in theory, impact both the function of the M protein and the topology of the M-related protein, we compared fibrinogen and C4BP binding in the emergent compared to the historic strains. For fibrinogen binding, our preliminary data noted strain to strain variability, so we assayed more than one historic and emergent strain. As shown in Fig. 4A, we observed no statistically significant binding difference amongst the emergent (ABC208 and TSPY637), historic (ABC25 and ABC3), and ABC25-chimera strains (P > 0.05 by ANOVA). GAS uses M protein, including M4, to bind C4BP (9,29,48) thereby inhibiting complement activation and conferring phagocytosis resistance, which could account for the increased survival in human blood previously identified for the emergent strains [54, 55]. By flow cytometry analysis, we identified no significant difference in binding of C4BP amongst strains ABC25, ABC208, and ABC25-chimera (P > 0.05 by ANOVA) (Fig. 4B). Consistent with M protein being the major mechanism by which GAS binds C4BP, we identified no significant C4BP binding to strain ABC208Δ*emm* relative to the negative control (Fig. 4B).

**FIG 4.**
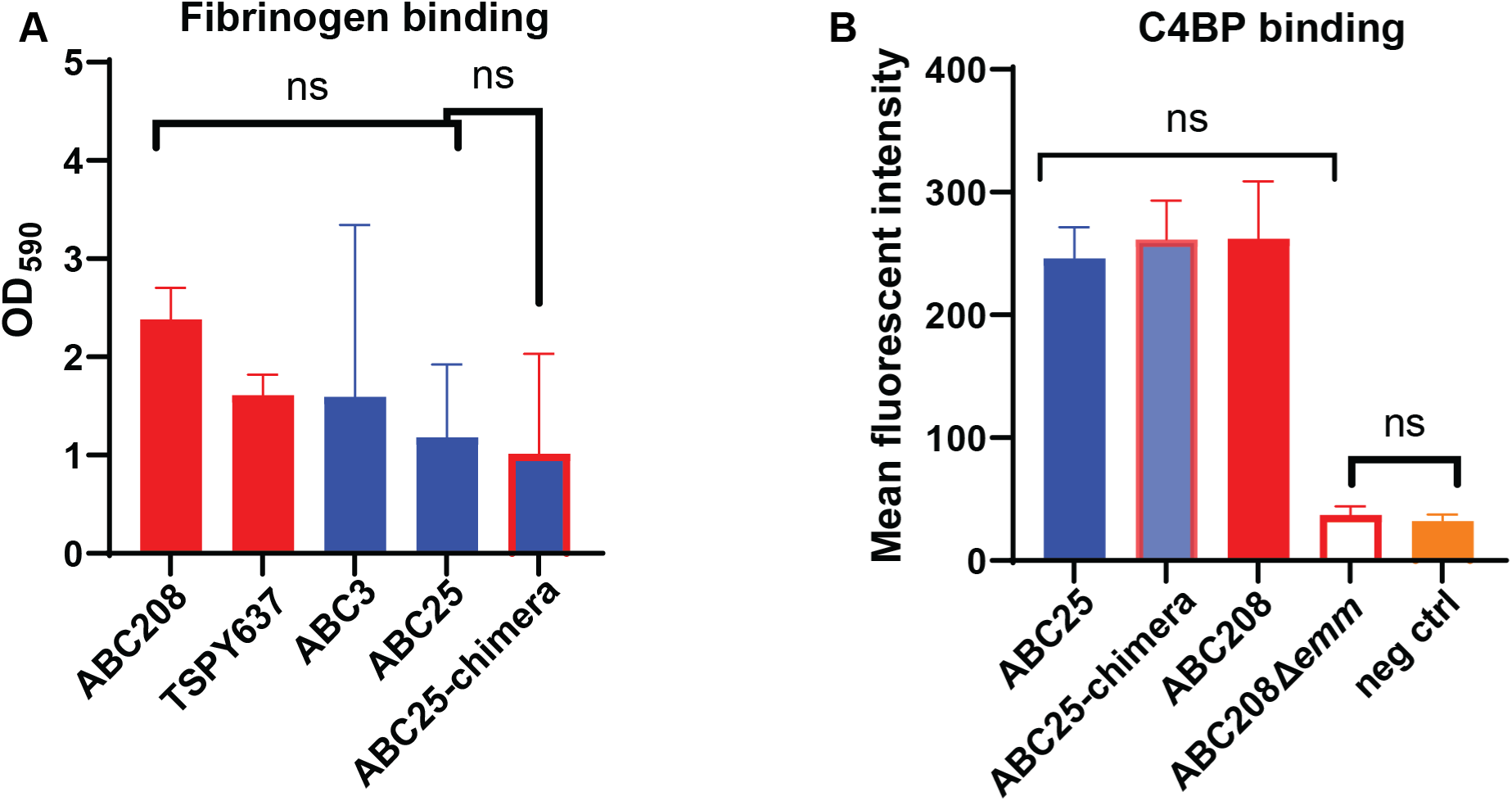
Analysis of interaction between the historic and emergent *emm4* strains and immune proteins. (A) Fibrinogen binding to the emergent (red), historic (blue), and historic strain with emergent M protein (blue and red). (B) C4BP binding to the emergent (red), historic (blue), and historic with emergent M protein (blue and red). The error bars represent the standard deviations for two replicates per strain on three separate days. NS indicates *P* > 0.05 by either ANOVA (three or more strain comparison) or Student’s t-test (two strain comparison).

### Identification of distinct GAS-macrophage interaction for emergent vs historic *emm4* GAS

Our previous study showed that the emergent strains survived better in non-immune human blood compared to the historic strains [31]. Thus, we next sought to test the hypothesis that the *emm*-*enn* chimera contributed to this phenotype by comparing the survival of the historic, historic-chimera, and emergent strains in macrophages. A significantly higher percentage of the original ABC208 inoculum survived intracellularly after 2 hours of macrophage interaction relative to ABC25 (P < 0.05), but there was no significant survival difference between ABC25 and ABC25-chimera (Fig. 5A) (*P* > 0.05). Thus, contrary to our hypothesis, the chimera *emm-enn* gene does not seem to mediate the differences in interaction with host immune cells observed between historic and emergent strains. To determine whether there are differences in strain interaction with other critical parts of the immune system, we repeated the analysis using polymorphonuclear leukocytes (PMNs or neutrophils). Similar to what was observed with the macrophages, the emergent strain ABC208 survived better in human neutrophils than the historic strain ABC25 (*P* < 0.05) (Fig. 5B).

**FIG 5.**
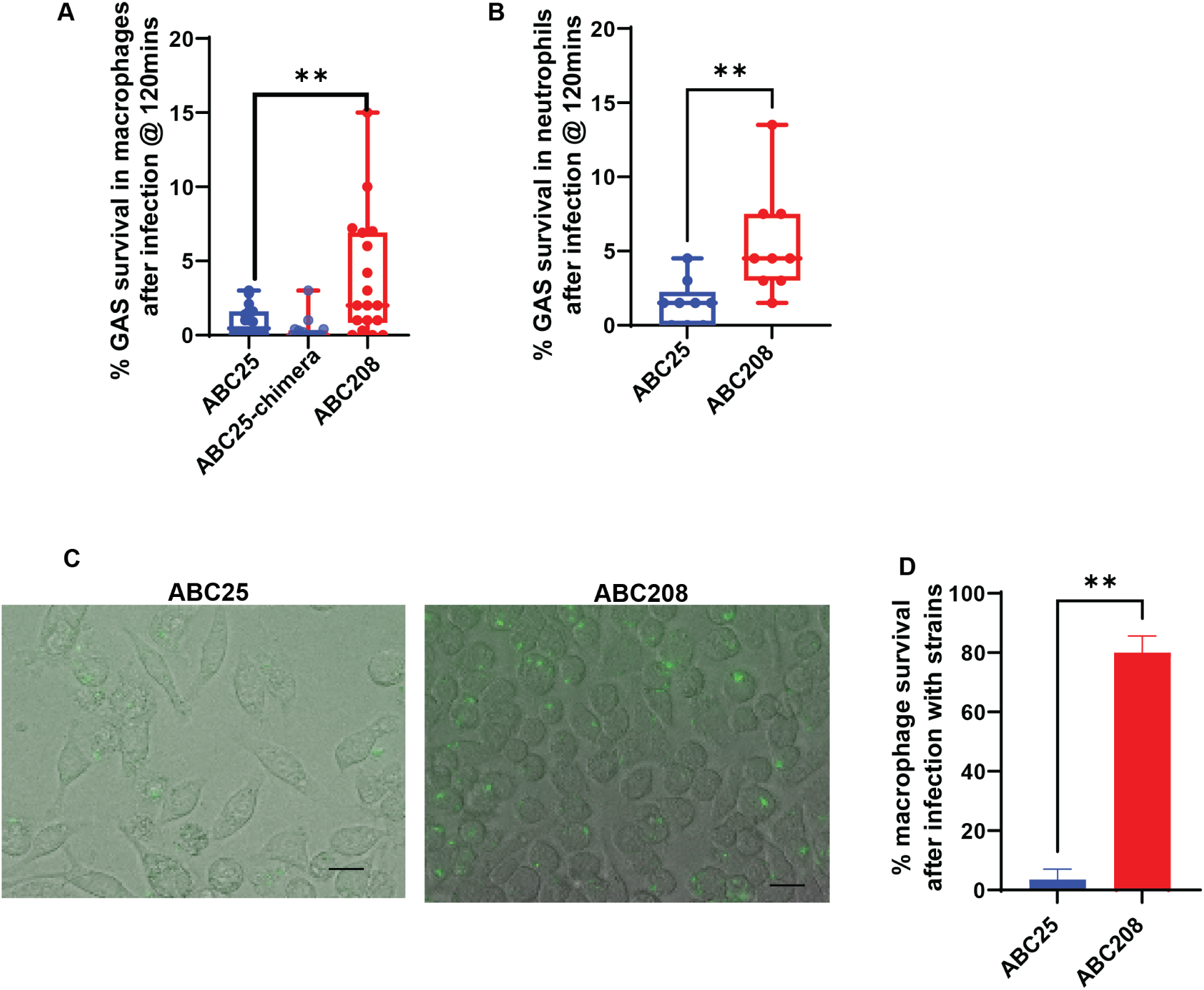
Analysis of interaction of historic and emergent *emm4* strains with immune effector cells. (A) Percent survival of an historic (blue), an emergent (red) and an historic strain with an emergent M protein (blue and red) following interaction with J774A.1 macrophages. (B) Percent survival of an historic (blue) and an emergent (red) strain following interaction with human neutrophils. (C) Representative section of light microscopic analysis of macrophages 12h post-infection with indicated GAS strains expressing GFP. (D) Viable macrophage analysis using trypan blue. The error bars denote the standard deviation of three biological and two technical replicates. For panels A, B, and D, ** = P < 0.05 by Student’s t-test).

To gain further insight into the differences in macrophage interaction between the emergent and historic strains, we visually analyzed GAS-macrophage interaction by infecting macrophages with GFP-tagged ABC25 and ABC208. Via light microscopy, we observed severe damage to the macrophages infected with ABC25 compared to ABC208 (Fig. 5C). Subsequent quantification with trypan blue showed about 20% of the macrophages infected with ABC25 were alive compared to about 80% for the macrophages infected with ABC208 (*P* < 0.05) (Fig. 5D). Consistent with the augmented recovery of ABC208 following macrophage lysis (Fig. 5A), we observed numerous instances of GFP expressing ABC208 inside of macrophages after 12 hours (Fig. 5C). Thus, these data show a marked difference in interaction between the emergent and historic *emm4* strains with macrophages.

### Identification of the critical role of Slo in macrophage killing by historic *emm4* GAS

The pore-forming toxin Slo has been identified as necessary and sufficient for inducing apoptosis in macrophages [56], and our previous study [31] showed that the historic strains have higher *nga*/*slo* transcript levels compared to the emergent strains. Thus, we next sought to determine whether Slo played a role in the observed macrophage damage by creating and assessing a historical strain lacking *slo*, ABC25Δ*slo*. The absence of Slo activity in ABC25Δ*slo* was confirmed by Slo hemolysis assay (Fig. 6A). We then used confocal microscopy in combination with the 7-AAD live/dead stain to analyze macrophages after infection with ABC25, ABC208, and ABC25Δ*slo*. By visual inspection, deleting *slo* abrogated macrophage damage by ABC25 to a level similar to that of ABC208 (Fig. 6B). We quantitated the macrophage death using Imaris imaging analysis and found ABC25 killed a higher percentage of macrophages compared to either ABC208 or ABC25Δ*slo* (Fig. 6C).

**FIG 6.**
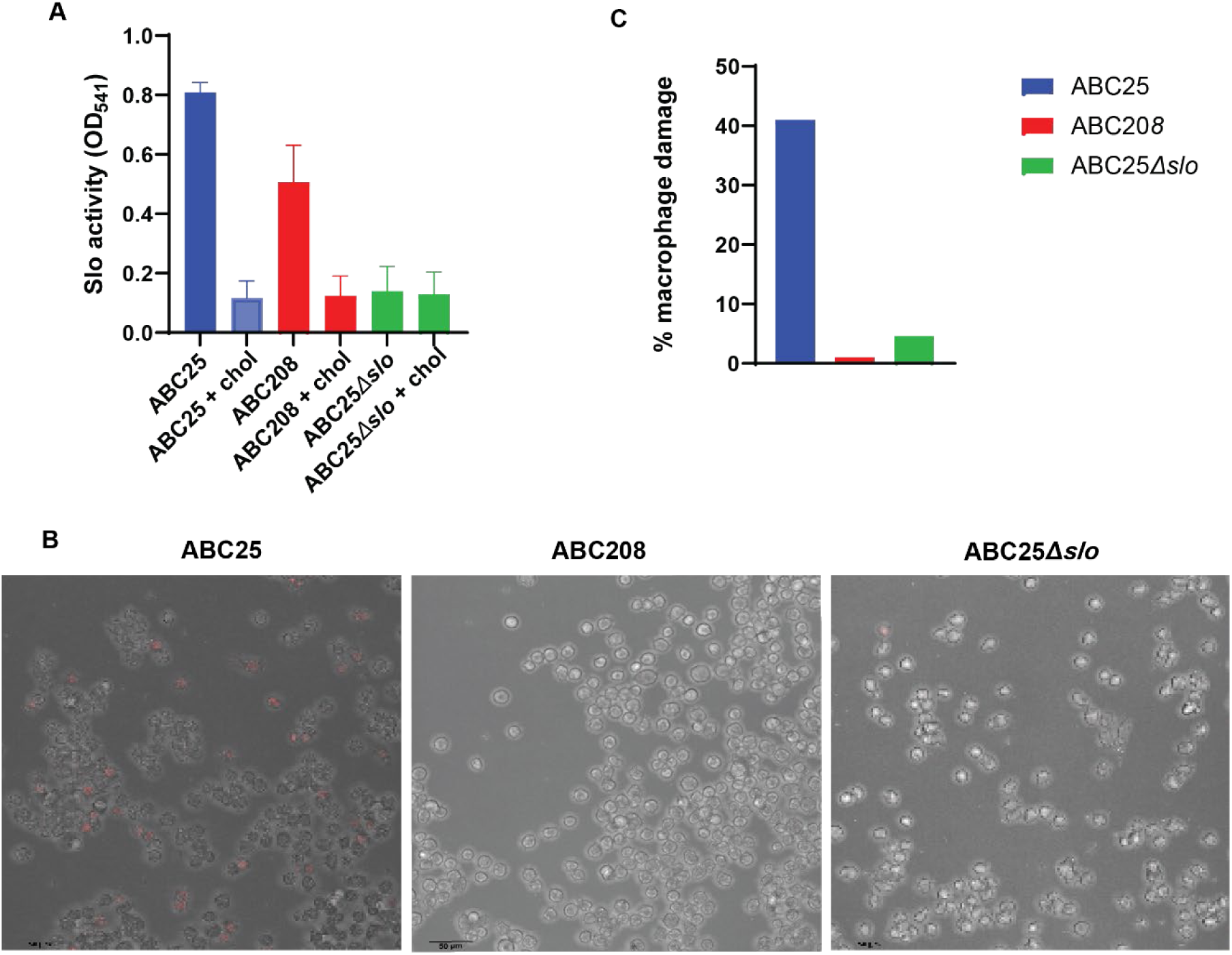
Microscopic analysis of macrophage killing by *emm4* GAS strains. (A) In vitro hemolysis assay to assess Slo activity (chol represents cholesterol in the control samples to inhibit Slo). (B) Representative sections of confocal microscopy of macrophages 3 hrs post-infection infected with indicated strains. Red staining by 7-AAD dye indicates dead cells. (C) Percent of dead macrophages following 3 hr incubation with indicated strains as identified by 7-AAD dye and quantified using Imaris imaging software.

### Extracellular and intracellular oxidative stress survival analysis in historic vs. emergent strains

Inasmuch as there was improved survival of the emergent strains during macrophage and PMN interaction (Fig. 5A/B), we reanalyzed our previously published RNAseq data comparing historic vs. emergent strain for possible “survival” strategies [31]. We noted augmented transcript levels in the emergent strains of the *msrAB* operon, which has been previously identified as involved in extracellular oxidative stress in various bacterial strains [57, 58] (Fig. 7A, 7B). This finding led us to test the hypothesis that the emergent strains had improved resistance to extracellular oxidative stress that GAS would encounter during interaction with macrophages and PMNs. We found that the emergent strains ABC208, TSPY637, and ABC199 survived significantly better in H_2_O_2_ than the historic strains ABC25, ABC3, and ABC221 (P < 0.05) (Fig. 7C). We further tested resistance to oxidative stress using sodium hypochlorite and again found that the emergent strains survived at a significantly higher level relative to the historic strains (P < 0.05) (Fig. 7D).

**FIG 7.**
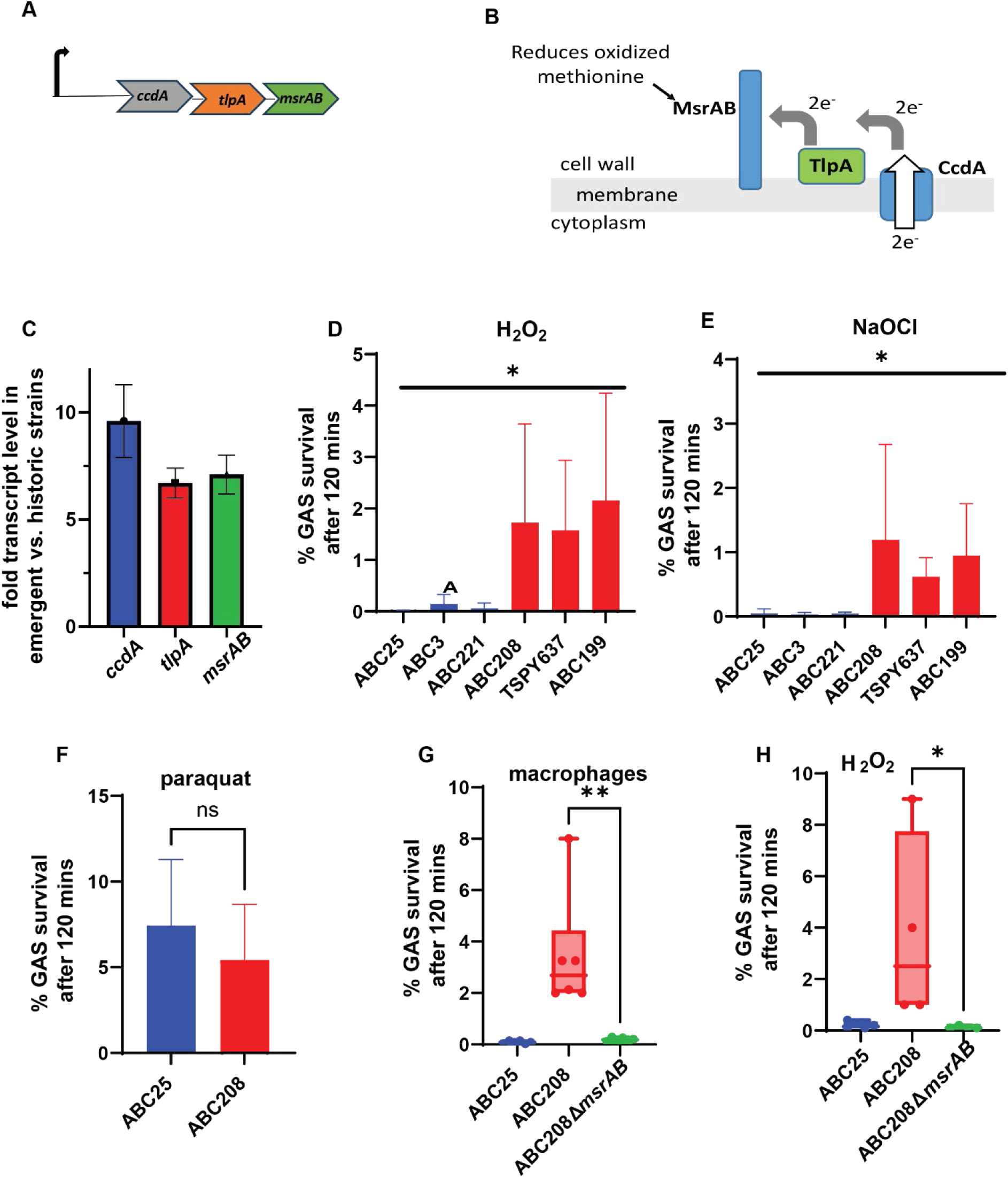
Extracellular and intracellular oxidative stress analysis. (A) Schematic of *ccdA-tlpA-msrAB* operon and (B) predicted protein location and function. (C) RNAseq analysis of c*cd-tlpA-msrAB* operon of four emergent relative to four historic strains as previously noted in [31] (D) Survival of the historic strains (blue) and the emergent strains (red) following exposure to H_2_O_2_. (E) Survival of the historic strains (blue) and the emergent strains (red) in the presence of NaOCl. For panels D and E, * indicates *P* < 0.05 by ANOVA. (F) Survival of the historic (blue) and the emergent (red) strains in the presence of paraquat. (G) Survival of strains ABC25 (blue), ABC208 (red), and ABC208Δ*msrAB* (green) following interaction with macrophages and (H) in the presence of H_2_O_2_. The error bars represent the standard deviation of three biological and two technical replicates. For panels F, G, and H, NS or * refers to P values > 0.05 and < 0.05, respectively, as determined by Student’s t-test.

In *Streptococcus pneumoniae*, the *msrAB* operon encodes proteins that are surface localized and shown to be specifically involved in response to extracellular but not intracellular oxidative stress [59–61]. Therefore, to determine whether the emergent strains were specifically resistant to extracellular oxidative stress, the strains were exposed to paraquat which induces the formation of reactive oxygen species (ROS), such as superoxide, in the cytoplasm in the presence of oxygen [60]. We identified no significant difference in the survival of ABC25 and ABC208 following paraquat exposure (Fig. 7E) (*P* > 0.05), consistent with a specific advantage of the emergent strains in withstanding extracellular oxidative stress.

To assess the role of *msrAB* in the increased oxidative stress resistance displayed by emergent strains, *msrAB* was deleted in ABC208 to create strain ABC208Δ*msrAB*. In accord with the critical role of MsrAB in the emergent strain defense against extracellular oxidative stress, ABC208Δ*msrAB* showed significantly reduced intracellular survival in macrophages (Fig. 7F) and reduced resistance to H_2_O_2_ to levels that were typically observed for ABC25 (P < 0.05 relative to parental strain ABC208) (Fig. 7G).

## DISCUSSION

Emergence and regression of microbial pathogens have long been major driving forces in human health [62, 63]. Even in the fairly short time since the delineation of GAS in the early part of the 20^th^ century [64] , numerous oscillations in GAS infections have been clearly recognized, such as the near disappearance of rheumatic fever in the United States and elsewhere over the past 50 years [65] contrasted by a surge of lethal *emm1* GAS disease since the 1980s [66]. Herein, we sought to unravel the mechanisms behind our recently discovered clonal replacement in *emm4* GAS in the United States and United Kingdom that occurred between 1995 and 2015 [29, 31]. In contrast to previous studies that correlated GAS clonal emergence with either acquisition of or augmented production of virulence factors [25, 67–70], we found that the emergent *emm4* GAS strains have a “survival” phenotype during interaction with immune cells relative to the historic strains, which readily killed macrophages. We correlated this phenotype with augmented defense against extracellular oxidative stress likely mediated by upregulation of the *msrAB* operon along with decreased production of the streptolysin O toxin.

One of the major genetic changes that delineates the emergent from historic *emm4* strains is the recombination between the *emm* and *enn* genes which resulted in a chimeric M protein [29, 31]. Given that M protein is the GAS major virulence factor and that the recombination event was universally present in emergent *emm4* strains [71], we initially hypothesized that the chimera was a major driver In the *emm4* replacement event. In a systematic study of M proteins from various *emm* types, the *emm4* protein was found to bind C4BP through an N-terminal site which is conserved between emergent and historic strains [72]. Additionally, the C-terminus is considered important for the coiled-coiled nature of the M protein such that variation in the C-terminus could impact overall M protein function [73]. Courtney et al. previously showed that *emm4* GAS strains bind fibrinogen and that this binding was important for growth of *emm4* GAS in human blood, which we previously observed was augmented in emergent GAS strains [52]. In contrast to our hypothesis, we observed no significant difference in the binding of a diverse array of host components to the emergent vs. historic strains nor did introducing the chimera M protein into a historic strain alter its interaction with macrophages. Since our original description of the chimera (65), recombination events in M and M-like proteins have been reported as isolated events in other M types [74]. These data suggest that the chimera has occurred on multiple occasions, but, other than in *emm4*, has not become fixed in a particular population, consistent with our finding of a lack of functional impact of the *emm-enn* chimera on *emm4* GAS pathophysiology. Thus, at present, we believe that the *emm-enn* fusion may be more of a “carrier” rather than “driver” genetic variation in the *emm4* population although we cannot entirely exclude the possibility that the *emm*/*enn* fusion may confer an advantage we heretofore have not uncovered.

A key finding of our work was the identification that emergent *emm4* GAS adopts a “survival” strategy during interaction with host immune cells in contrast to the destructive phenotype evidenced by the historic strains. We linked this finding with increased tolerance of the emergent strains to extracellular oxidative stress, possibly due to increased expression of the *msrAB* operon. This “survival strategy” could assist GAS by at least two major mechanisms. First, GAS faces a fairly high concentration of oxygen in the upper airway and lacks the catalase enzyme that many organisms use to detoxify H_2_O_2_ [75]. Additionally, ROS generated by phagocytotic cells such as macrophages and neutrophils during the innate immune response is a major mechanism for GAS clearance, and the GAS defense against ROS has been shown to be important to pharyngeal colonization in diverse animal models [57, 75]. MsrAB, in conjunction with its two accessory components, TlpA and CcdA, contributes to oxidative stress defense by combating oxidation of methionine residues which generates methionine sulfoxide, which in turn can alter protein function through confirmational changes [57, 58, 76]. Thus, emergent *emm4* GAS strains may have an advantage in directly surviving the oxidative stress encountered during host-pathogen interaction. A second major mechanism by which the “survival strategy” could assist *emm4* GAS is by turning macrophages into a “trojan horse” for the emergent strains [77]. Epithelial and macrophage-like cells have been shown to be reservoirs for intracellular GAS, which are associated with recurrent pharyngotonsillitis [78, 79]. By not destroying but rather surviving inside macrophages, the emergent strains may be evading key aspects of the host immune response, thereby gaining an additional competitive advantage [80]. Intriguingly, the emergent strains have a growth defect in laboratory conditions which appears to result from a conserved mutation in the gene encoding carbonic anhydrase, a protein that catalyzes the conversion of CO_2_ and H_2_O to HCO3- and H+ [81]. Carbonic anhydrase has been proposed as a novel antimicrobial target [82], yet *emm4* GAS has remained one of the most common causes of GAS infection despite apparently lacking a functional enzyme [83]. Why emergent *emm4* GAS contain a conserved mutation that causes a fitness defect during growth in laboratory conditions is an intriguing question that our laboratory is currently exploring given that such fitness defects are typically considered an explanation for lack of infectivity by a microbe [84, 85].

Our finding of a correlation between a “survival” strategy and strain replacement in *emm4* GAS adds to the understanding of factors driving GAS clonal emergence. Using *emm1*, *emm3*, and *emm89* as models, large-scale studies of recently circulating GAS strains, often primarily collected from invasive infections, have consistently correlated strain prevalence with acquisition or augmented production of virulence factors [18, 66, 68]. Yet GAS has been co-evolving with humans for thousands of years [86] and a cycle of ever-increasing virulence is highly unlikely to be conducive to long-term GAS evolutionary benefit [87]. We hypothesize that the “survival strategy” of the recent *emm4* GAS strains may have been silently occurring in other *emm* types, which could contribute to the emergence and then near disappearance of serious GAS infections, such as that occurred with scarlet fever in the early 1900s [88]. It is potentially noteworthy that recent systematic studies of *emm4* GAS have consistently found that *emm4* strains are among the most common causes of pharyngitis but have a very low “invasive” index meaning that the number of invasive infections is lower than expected relative to pharyngeal cases [89, 90]. It was previously found that carbonic anhydrase from *S. pneumoniae* is critical in animal models where the bacterium needs to cross epithelial and endothelial cell boundaries for dissemination into the bloodstream [49]. Therefore, the carbonic anhydrase mutation in the emergent strains, along with low levels of virulence factor production, may contribute to the low *emm4* invasive index. Systematic studies of non-invasive, in addition to invasive, GAS isolates may further augment understanding of the highly impactful oscillations of GAS disease in humans.

In conclusion, we have extensively analyzed the genetic and phenotypic changes separating emergent from historic *emm4* GAS strains. Rather than a single event leading to acquisition or heightened production of a virulence factor as has previously been identified in GAS clonal emergent events, we identified the development of a “survival” strategy by the emergent strains during interaction with key components of the innate immune system. These data augment understanding of GAS clonal emergence strategies, which are likely applicable to a broad array of major human pathogens.

## MATERIALS AND METHODS

### Bacterial strains and growth conditions

The bacterial strains used in this study are shown in Table 1. Representative historic and emergent *emm4* strains were selected from our collection of >1,000 isolates with whole genome sequencing data based on their representing major sub-clades and lacking mutations in major regulatory genes such as the control of virulence regulatory two-component system (*covRS*) [91]. GAS growth was performed without agitation in Todd-Hewitt broth with 0.2% yeast extract (THY), on THY agar, and on Trypticase soy agar with 5% sheep blood. For quantitative growth assessment assays, bicarbonate was added as previously described [92], growth was monitored in a BioTek Synergy H1 plate reader, and readings were obtained every 30 min following 10 seconds of shaking. When needed, antibiotics were added in the following concentrations (ampicillin at 100 μg/ml, spectinomycin at 150μg/ml, kanamycin at 150 μg/mL, chloramphenicol at 25 μg/ml).

**Table 1.** Strains used in this study.

| Strain | Description | Source |
| --- | --- | --- |
| ABC3 | Historic clade invasive *emm4* strain isolated in 1997 in the United States | [31] |
| ABC25 | Historic clade invasive *emm4* strain isolated in 1997 in the United States | [31] |
| Duke | Emergent clade invasive *emm4* strain isolated in 2014 in the United States. The source for *emm-enn* gene fusion cloned into strain ABC25 | [108] |
| ABC221 | Historic clade invasive *emm4* strain isolated in 2006 in the United States | [31] |
| ABC199 | Emergent clade invasive *emm4* strain isolated in 2005 in the United States | [31] |
| ABC208 | Emergent clade invasive *emm4* strain isolated in 2005 in the United States | [31] |
| TSPY637 | Emergent clade invasive *emm4* strain isolated in 2015 in the United States | [31] |
| ABC25-chimera | ABC25 with emergent *emm/enn* from *emm4* strain Duke | This study |
| ABC25Δ*slo* | ABC25 with complete removal of *slo* gene | This study |
| ABC25*saca*Q12* | ABC25 with emergent *saca* gene | This study |
| ABC208Δ*emm* | ABC208 with complete removal of *emm* gene | This study |
| ABC208Δ*msrAB* | ABC208 with complete removal of *msrAB* gene | This study |
| ABC208*ralp3*<sup>WT</sup> | ABC208 with restored wildtype *ralp3* gene | This study |
| ABC208*saca*<sup>WT</sup> | ABC208 with restored wildtype *saca* gene | This study |
| ABC25-GFP | ABC25 with GFP expressing plasmid | This study |
| ABC208-GFP | ABC208 with GFP expressing plasmid | This study |

### DNA manipulation

The plasmids and primers utilized in this work are shown in Table 2. Plasmid DNA was extracted using QIAprep Spin Miniprep Kit (Qiagen, Germantown MD USA) and used to transform *E. coli* and GAS strains by electroporation, as previously outlined [48]. The isogenic mutant strains, ABC208Δ*emm,* ABC25Δ*slo,* and ABC208Δ*msrAB* were created by non-polar insertional mutagenesis with a spectinomycin resistance cassette as previously described [93]. Homologous recombination was used to construct ABC25-chimera by substituting *emm* with the *emm/enn* fusion gene from the emergent *emm4* strain Duke [29] as previously described in [94]. Isoallelic exchange was used to construct ABC208*ralp3^WT^* by replacing the truncated *ralp3* with the full-length *ralp3^WT^* in ABC208 as previously described [95]. ABC208*saca^WT^* and ABC25*sacaQ12\** were constructed by employing a counterselection technique using levansucrase (*sacB*) maker as described in [96] with some modifications. SacB secretes levansucrase which is lethal to GAS in a high sucrose environment. PCR was used to amplify wildtype (*saca*^WT^) *saca* from ABC25 and *saca* mutant (*saca*Q12*) from ABC208. Both amplified segments were ligated separately into the pJL1055 plasmid containing *sacB* and chloramphenicol cassette. Both clones were electroporated into ABC208 and ABC25 respectively. The clones were subsequently passaged without chloramphenicol and plated in THY Agar containing sucrose to select for the mutants ABC208*sacA^WT^*and ABC25*sacAQ12\** respectively. Gfp-tagged strains ABC25 and ABC208 were constructed by amplifying the superfolder GFP (sfGFP) cassette from pET28-sfGFP using the listed primers (Table 2). The Cfb promoter (pCfb) from group B *Streptococcus* were amplified for fusion to the GFP cassette to drive constitutive fluorescent protein expression in GAS as detailed in [97]. pCfb and sfGFP fragments were fused by splicing overlap extension (SOEing) PCR using primers as listed, and the resulting product was introduced by restriction digest into the multicopy self-replicating plasmid pLZ12Km2 [98] to generate pLZ12Km2: sfGFP. The pLZ12Km2: sfGFP plasmid was maintained in GAS by culturing in the presence of Kanamycin (150µg/ml).

**Table 2.**
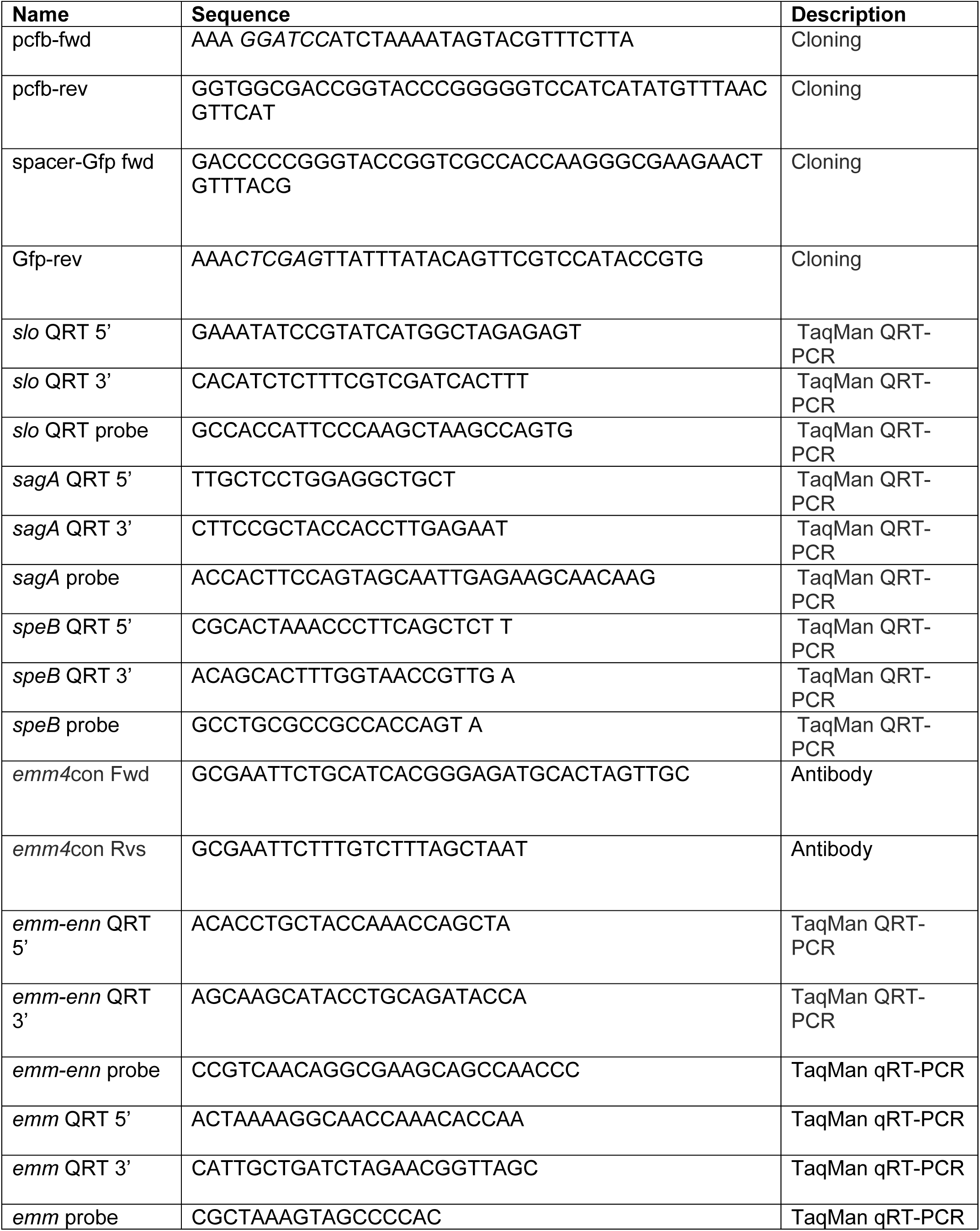
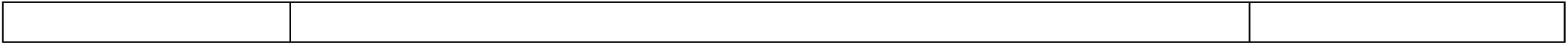
Primers and probes.

### Evaluation of the Sil system

A 17 amino acid mature SilCR peptide (DIFKLVIDHISMKARKK) was synthesized by Genscript. After reconstitution, different concentrations were added to the cell culture as previously described [99]. For SilCR-induced gene expression analysis, overnight cultures of two historic (ABC25 and ABC3 and two emergent (ABC208 and TSPY637) *emm4* strains were added to prewarmed THY medium containing SilCR at concentrations of 0, 10, or 500 ng/mL SilCR. The cultures were incubated at 37 °C under 5% CO_2_ and the bacteria were collected during the mid exponential phases followed by qRT-PCR. The experiment was performed three times in duplicate.

### Transcript level analysis

Transcript level analysis was performed using TaqMan qRT-PCR on strains grown to the mid-exponential (ME) phase in THY medium and analyzed using an Applied Biosystems Step-One Plus System as described [100]. Primers and probes are listed in Table 2. Experiments were performed at least in duplicate on two separate days.

### Assessment of M protein production and cell surface expression

For total M protein production assessment, GAS strains were grown to ME phase in 45ml THY. Cell cultures were centrifuged, and pellets were resuspended in lysis buffer (1M Tris, 0.5M EDTA, Triton X-100, mutanolysin and lysozyme) for 10 minutes. Protein concentration was determined using the Bradford assay, and equal amounts of total protein were then separated using sodium dodecyl sulfate-polyacrylamide gel electrophoresis (SDS-PAGE). M protein was detected using a custom-made anti-M4 antibody (Covance) raised in rabbits against the N-terminal domain of the mature M4 protein (amino acids 42-165) and visualized on the Odyssey imaging system using a goat anti-rabbit secondary antibody (LI-COR Biosciences) as described previously [101]. For cell surface detection of M protein, strains were grown in THY medium to exponential phase, harvested, washed with phosphate-buffered saline (PBS), and resuspended at 10^8^ CFU/ml. Anti-M4 protein antibody was added at a final concentration of 0.05ug/ml in a total volume of 100µl and incubated on ice for 30 minutes. Samples were washed with veronal buffer and incubated with Alexa Fluor 647 Affinity Pure Goat Anti-Rabbit IgG (H+L) on ice for 30 minutes. Cells incubated in only primary and only secondary antibodies were used as controls. The samples were washed with a staining buffer and analyzed using flow cytometry (BD LSRFortessa).

### Transmission electron microscopy

Cells were exposed to the radiofrequency field for 5 minutes and fixed using a solution of 3% glutaraldehyde and 2% paraformaldehyde in 0.1 M cacodylate buffer at pH 7.3. Cells were then washed in 0.1 M sodium cacodylate buffer, treated with 0.1% Millipore-filtered cacodylate buffered tannic acid, post-fixed with 1% buffered osmium, and stained altogether with 1% Millipore-filtered uranyl acetate. The samples were dehydrated using increasing concentrations of ethanol, then infiltrated and embedded in LX-112 media. The samples were polymerized in a 60°C oven for 3 days. Ultrathin sections were cut with a Leica Ultracut microtome (Leica, Deerfield, IL), stained with uranyl acetate and lead citrate using a Leica EM Stainer, and examined with a JEM 1010 transmission electron microscope at 80 kV. The digital images were obtained using the AMT Imaging System (Advanced Microscopy Techniques, Danvers, MA).

### GAS-fibrinogen binding assay

Comparison of fibrinogen between the historic and the emergent strains was performed as described with some adjustments [102]. An overnight culture of the strains was added to prewarmed THY medium, grown to ME phase, and serially diluted. The serially diluted bacteria were added to the wells of microtiter plates that had been coated with fibrinogen (10 μg/well). Adhered bacteria were treated with 4% formaldehyde and then stained with 0.5% crystal violet. Following washing, 50 µl of a 10% acetic acid solution was added and the absorbance was measured at 590 nm.

### GAS cell surface assessment of C4 binding protein (C4BP)

Deposition of C4BP on the GAS cell surface was performed as described in [103] with slight modifications. Strains were grown to ME phase, collected by centrifugation, and washed cells were incubated with 10% normal human serum (Complement Technology Inc) for 30 minutes Bacteria were then washed twice with veronal buffer and resuspended in staining buffer containing anti-C4BP monoclonal antibody (A215, Quidel) on ice for 30 minutes. Cells were washed twice with veronal buffer and stained with secondary antibodies coupled to PE. Finally, the bacteria were washed with veronal buffer and resuspended in 500 μl of sorting buffer for flow cytometry analysis using a Cyflow space flow cytometer (BD LSRFortessa). The experiment was repeated twice in duplicate.

### Assessment of GAS survival inside macrophages

These assays were performed as previously described in [56] with some modifications. J774A.1 macrophages (ATCC TIB-67) were cultured in Roswell Park Memorial Institute (RPMI) 1640 Media with 10% fetal bovine serum (FBS) in the presence of penicillin and spectinomycin (10mg/ml) and seeded at 5x10^5^ cells per well in a 24-well plate a day prior to the assessment. Macrophages were infected with GAS strains at a multiplicity of infection (MOI) of 10 in 350µl RPMI with 10% fetal bovine serum FBS without antibiotics. The plate was centrifuged at 2000 rpm then incubated for 30 minutes before penicillin (5 µg/ml) and gentamicin (100 µg/ml) were added to each well for an additional 45 minutes to kill extracellular bacteria, and subsequently washed with PBS. At different time points following antibiotic removal, macrophages were lysed with 0.1% Triton, serially diluted, and the dilutions were plated to quantify the CFU. The % survival was calculated as (CFU recovered at 120 minutes /input CFU) x 100.

### Direct visualization of GAS-macrophage interaction

Macrophages were infected with GFP-tagged ABC25 and ABC208 and incubated for 30 minutes. The extracellular bacteria were killed as described above. After washing, the interaction between the macrophages and the GFP-tagged bacteria was analyzed using light microscopy (Keyence BZ-X800) over 12 hours. To analyze the impact of SLO on GAS-macrophage interaction, we were unable to use GFP due to antibiotic selection incompatibility between the GFP plasmid and the Slo knockout strain. Thus, we used confocal microscopy (Leica SP8 Laser Scanning, Leica Microsystems) with the 7-AAD live/dead stain (Thermo Fisher) [104] with experiment performed over 3 hours. Quantitation of macrophage viability was done with the Imaris imaging analysis software (Oxford Instruments).

### Macrophage quantification with trypan blue exclusion test for cell viability

Trypan blue dye exclusion test was used to assess cell integrity as described [105]. Macrophages were infected at MOI of 10 and incubated for 1 hr. After incubation, the cells were scraped from the bottom of the flask, dyed with trypan blue, and quantified using a cell counter (Invitrogen™ Countess™ 3 Automated Cell Counter) to analyze dead and live macrophages.

### SLO enzyme-activity assay

The amount of secreted Slo was assessed as described in [106] with some modifications. Samples were grown to OD_600_ of 0.45. The cells were centrifuged, and the supernatant was filtered with a 0.22 μm pore size filter. 20 mmol/L dithiothreitol was added to the filtered supernatant and incubated at room temperature for 10 min. The incubated samples were aliquoted into 2 tubes (500 μL each) and water-soluble cholesterol (25 μg) (cholesterol/methyl-β-cyclodextrin; Sigma-Aldrich) was added to only one of the samples and both samples were incubated at 37°C for 30 min. 250 μL of 2% sheep erythrocyte/PBS suspension was added to each sample, mixed by inversion, and incubated for 30 min at 37°C. 500 μL of PBS was added to each sample and subsequently centrifugated at 3000 *g* for 5 min to separate the unlysed erythrocytes. The amount of hemoglobin present in the supernatant was measured in the spectrophotometer at OD 541 nm. The positive control was erythrocytes incubated in water, and fresh THY broth was used as a negative control. The experiment was repeated twice in duplicate.

### Isolation of Human PMNs and GAS survival assay

Human polymorphonuclear neutrophils (PMNs) were isolated from the venous blood of healthy individuals following informed consent using a STEMcell isolation kit (STEMCELL Technologies). The purified neutrophils were resuspended in RPMI medium. Neutrophil viability was assessed via trypan blue (Sigma-Aldrich, St. Louis, MO, USA) prior to the experiment. Neutrophils (5x10^6^) were seeded in 24-well plates at MOI of 10 and incubated at 37°C in 5% CO_2_ for total of 120 minutes and antibiotics were used to kill the extracellular bacterial cells. At different time points, the wells were washed with PBS, and cells were centrifuged at 300 rpm for 20 minutes before lysing with saponin. Recovered bacterial cells were serially diluted, plated on blood agar plates, incubated overnight and the CFU was enumerated for survival analysis.

### Oxidative stress assays

Resistance to killing by hydrogen peroxide (H_2_O_2_), sodium hypochlorite (NaOCl), and paraquat was assayed as described in [57, 107] with some modifications. Strains were grown to 0.5-0.7 OD_600_), and oxidative stress was added (H_2_O_2_ 12 nM; NaOCl 5 nM, and paraquat 40 nM). THY alone was used as control. Bacteria were enumerated after 2 hours of incubation, and the percent survival was calculated based on CFUs with and without the oxidative stress. The assay was repeated three times.

### Statistical analysis

GraphPad/Prism (v10) was used to perform all statistical analyses. Mann-Whitney U test was used for nonnormally distributed data while Student’s t test was used for comparing normally distributed continuous variables amongst two strains/conditions whereas ANOVA was used for three or more strains/conditions. A *P* value less than 0.05 was considered as statistically significant.

## ACKNOWLEDGEMENT

This work was supported by NIH R21AI142126 and R21AI159059 to S.A.S and A.R.F

